# Clutter does not affect the spatial tuning of category representations in human object-selective cortex

**DOI:** 10.64898/2026.07.28.741311

**Authors:** Karla Matić, Issam Tafech, Leonardo Pettini, Rony Hirschhorn, Kai Görgen, Carsten Bogler, Peter König, Radoslaw M. Cichy, John-Dylan Haynes

## Abstract

Natural vision is frequently impaired when the visual field is crowded with multiple competing objects. One common interpretation of this clutter interference is that it originates from the large, position-tolerant receptive fields found in high-level visual cortex, which hinder the isolation of a target from its spatial surround. One way the visual system might counteract this interference is by relying on more localized object representations that are less prone to interference from objects at other locations and that are known to be prevalent also in higher-level visual cortex. If object encoding under clutter indeed uses more localized representations, then classifiers trained to recognize objects from one region of the visual field should generalize less well to other regions of the visual field in the presence of clutter compared to isolated objects. Here we tested this by measuring participants’ brain signals with fMRI while they viewed objects presented either alone or within cluttered arrays. Consistent with prior work, category decoding accuracy was significantly reduced in clutter. However, we found that even classifiers trained directly on cluttered arrays generalize robustly across spatial positions, comparable to classifiers trained on isolated objects. This finding challenges the idea that clutter preferentially disrupts position-tolerant representations.

## Introduction

One of the striking features of natural vision is the efficiency with which we recognize objects (DiCarlo et al., 2012). Even a brief exposure to a visual stimulus triggers a cascade of neural activity along the visual stream, beginning with the processing of simple features in early visual areas (Carandini et al., 2005), and leading up to the encoding of complex object information in higher visual cortex (Bracci & Op de Beeck, 2016; Grill-Spector & Weiner, 2014; Hebart et al., 2020; Haxby et al., 2001; Kriegeskorte et al., 2008; Doerig et al., 2025).

Object recognition has mostly been studied using only single objects presented in isolation (e.g., Grill-Spector et al., 2001; Hebart et al., 2020; Kanwisher et al., 1997; Kriegeskorte et al., 2008; Op de Beeck et al., 2019). In natural vision, however, our visual field typically contains multiple visual inputs simultaneously, and the resulting clutter can degrade the recognition of each individual item (Rosenholtz et al., 2007; Bouma, 1970). This raises the question of how visual system deals with interference between multiple objects in the visual field (Li et al., 2009; Desimone & Duncan, 1995; Zoccolan et al., 2007; Reynolds et al., 1999; Chelazzi et al., 1998; Sundberg et al., 2009; Miller et al., 1993; MacEvoy & Epstein, 2009; Baeck et al., 2013; Cohen et al., 2014; Doostani et al., 2023; Cox & Riesenhuber, 2015; Reddy et al., 2009; Kastner et al., 1998; Kastner & Ungerleider, 2001; Bao & Tsao, 2018; Zoccolan et al., 2005).

In early visual regions, spatial interference by the surround, while clearly present (Allman et al., 1985), is limited in scope. This is due to the comparatively small receptive fields both at the single-cell and the population level (Dumoulin & Wandell, 2008; Smith et al., 2001; Klink et al., 2021), and the more local nature of surround interactions (Nurminen & Angelucci, 2014). However, higher-level visual areas have larger receptive fields, and interference even between spatially separated visual field locations is more prevalent (Decramer et al., 2019; Vanhoyland et al., 2026; Op de Beeck & Vogels, 2000; Kastner et al., 1998; Klink et al., 2021). This interference has been explained by various computational pooling mechanisms (Bao & Tsao, 2018; Rousselet et al., 2003; Zoccolan et al., 2005). One proposed solution to the problem of interference in higher-level visual areas is focused attention, which can provide a selection bias to individual target locations (Kastner & Ungerleider, 2001). In the absence of focused attention, other mechanisms might help sustain object representations in the presence of clutter, such as coarse coding in neural population codes (Chaney et al., 2014).

Another possibility, however, is that crowding in higher-level visual cortex might not be so problematic and interference-prone as thought. Although receptive field sizes increase strongly across the visual hierarchy in both monkeys (Klink et al., 2021) and humans (Larsson & Heeger, 2006), they are nevertheless highly variable: many neurons in higher visual areas still have relatively small receptive fields (Op de Beeck & Vogels, 2000), giving rise to considerable information regarding the spatial location of objects (Cichy et al., 2011; Sayres & Grill-Spector, 2008). Thus, it is conceivable that object encoding under clutter in high-level regions is performed by units with more localized receptive fields (so-called “position-dependent” as opposed to “position-tolerant” coding). This could be implemented either in a static code, potentially explaining why higher-level regions still have many units with smaller receptive fields, or in a dynamic shrinking of receptive fields to focus on local objects in the presence of clutter (Rolls, 2008). In both cases, the prediction is that object-related information should be more localized in cluttered conditions than in isolated displays.

Here we directly tested this hypothesis using human fMRI while participants observed single-object and cluttered visual displays. The aim was to assess whether object-related information in high-level object-selective cortex (LO) is more localized under conditions of clutter. To test this, we presented participants with either single-object or cluttered displays and assessed the object-related information using multivariate decoding (Haxby et al., 2014; Haynes, 2015; Haynes & Rees, 2006; Tong & Pratte, 2012). If indeed object-related information is more localized under clutter, we would expect a classifier trained under clutter to generalize worse to other locations than if trained for single-object displays (**Fig. 1A**).

**Figure 1.**
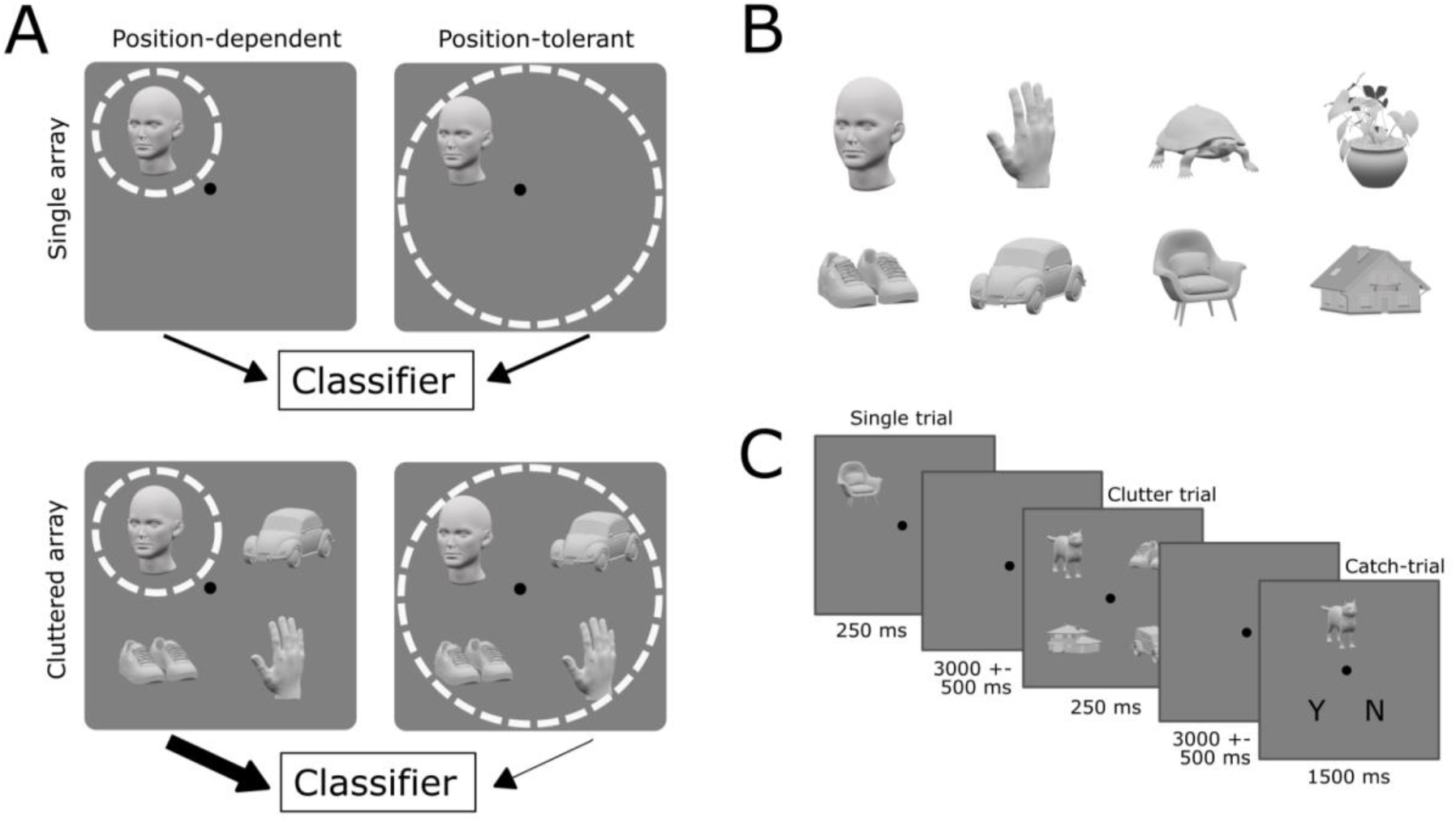
Experimental design. **A)** Position tolerance of object representations and decodable information. Area LO is known to have representational units that vary widely in their spatial tuning width (Op de Beeck & Vogels, 2000). *Position-dependent* representations (left column) are thought to arise from neurons with smaller receptive fields that only respond to an object when it is presented at a specific location. This also means they might be less sensitive to clutter in other regions of the visual field (bottom left). In contrast, *position-tolerant* representations (right column) involve larger receptive fields that are responsive to preferred objects at multiple locations, meaning they will also potentially be influenced by clutter at adjacent regions of the visual field (bottom right). In the absence of clutter (top row), position-dependent as well as position-tolerant units have information that can be used by a classifier to identify an object. In contrast, under clutter the object-related information that can be used by a classifier should be mostly restricted to position-dependent units (indicated by the thick arrow). This makes an important prediction: When trained for isolated objects, the position-tolerant representations included in the classifier will allow it to generalize across different visual field locations. In contrast, when training under clutter, the focus on position-dependent representations would mean a classifier should not generalize well to other spatial locations. **B)** Example objects from the stimulus set (for full details see Methods). Stimuli were selected from 8 categories: 4 animate (faces, hands, animals, plants) and 4 inanimate (shoes, cars, chairs, houses). For each category, 4 different images were shown: two different exemplars from the given category, each rotated around the vertical axis and presented in two orientations (45° left and 45° right), resulting in a set of 32 unique stimuli. **C)** Trial types in the fMRI experiment. Single-object and cluttered arrays were shown in random order for 250 ms each, followed by a blank inter-trial interval. In occasional catch trials (10% of trials), participants reported whether a probe object appeared in the preceding array.

## Results and Discussion

We tested whether, in the presence of clutter, object information in lateral occipital cortex is restricted to more localized, position-dependent representations (**Fig. 1A**). We used fMRI to measure brain activity in 19 participants while they viewed the same objects in isolation and in clutter (**Fig. 1B-C**). The clutter consisted of multiple isolated objects (Cox & Riesenhuber, 2015) in the four quadrants of the visual field (rather than objects superimposed on wide-field cluttered backgrounds as in e.g., Graumann et al., 2022). We applied linear classifiers to single-trial fMRI measurements to decode category information of each object. Most importantly, we performed generalization tests (see e.g., Cichy et al., 2011; Schwarzlose et al., 2008) to examine how category classifiers generalize across spatial positions and compared this between isolated and cluttered displays. If object information under clutter is focused more on position-dependent representations, this should manifest as a selective drop in cross-position generalization for classifiers trained in clutter.

In each participant, we defined the lateral-occipital cortex (LO), a region involved in high-level object recognition (Cichy et al., 2011; Grill-Spector et al., 2001; Paulun et al., 2025; see **Fig. 2A** and Methods for details). In line with previous studies (Bougou et al., 2024; Decramer et al., 2019; Vanhoyland et al., 2026), we focus on the lateral occipital object-selective cortex (LO) for several reasons. First, it is consistently informative about object identity in cluttered displays (Baeck et al., 2013; Graumann et al., 2022; MacEvoy & Epstein, 2009). Second, it broadly responds to various types of shapes and objects (Bougou et al., 2024; Decramer et al., 2019; Vanhoyland et al., 2026) rather than being highly specialized only for specific object categories. While interference between competing objects is reduced when each object in the visual field activates a dedicated region of cortex (Reddy & Kanwisher, 2007), in LO the potential interference should be maximized. Third, shape and object encoding in LO has been studied in detail also at the level of human single cells, thus allowing for comparison to the human fMRI signals measured here (Bougou et al., 2024; Decramer et al., 2019; Vanhoyland et al., 2026) and to monkey region IT (Bougou et al., 2024; Kriegeskorte et al., 2008). Finally, LO combines large receptive fields that include also ipsilateral and bilateral visual field representations on the one hand (Decramer et al., 2019; Klink et al., 2021; Vanhoyland et al., 2026; Op de Beeck & Vogels, 2000), with high variability in spatial extent (Op de Beeck & Vogels, 2000; Silson et al., 2016) and considerable location information on the other (Cichy et al., 2011; Sayres & Grill-Spector, 2008). Thus, it is uniquely suited for studying the potential role of large versus small receptive fields under clutter. To confirm that we did not miss out on any relevant information by focusing our analysis on the LO region, we additionally performed a post-hoc searchlight analysis identifying regions maximally informative about our object categories (**Fig. 2A**, blue region; see Methods for details). This analysis showed that the locus of category information largely overlaps with our independently defined ROI, extending slightly more anterior as has been previously observed (Kriegeskorte et al., 2008).

**Figure 2.**
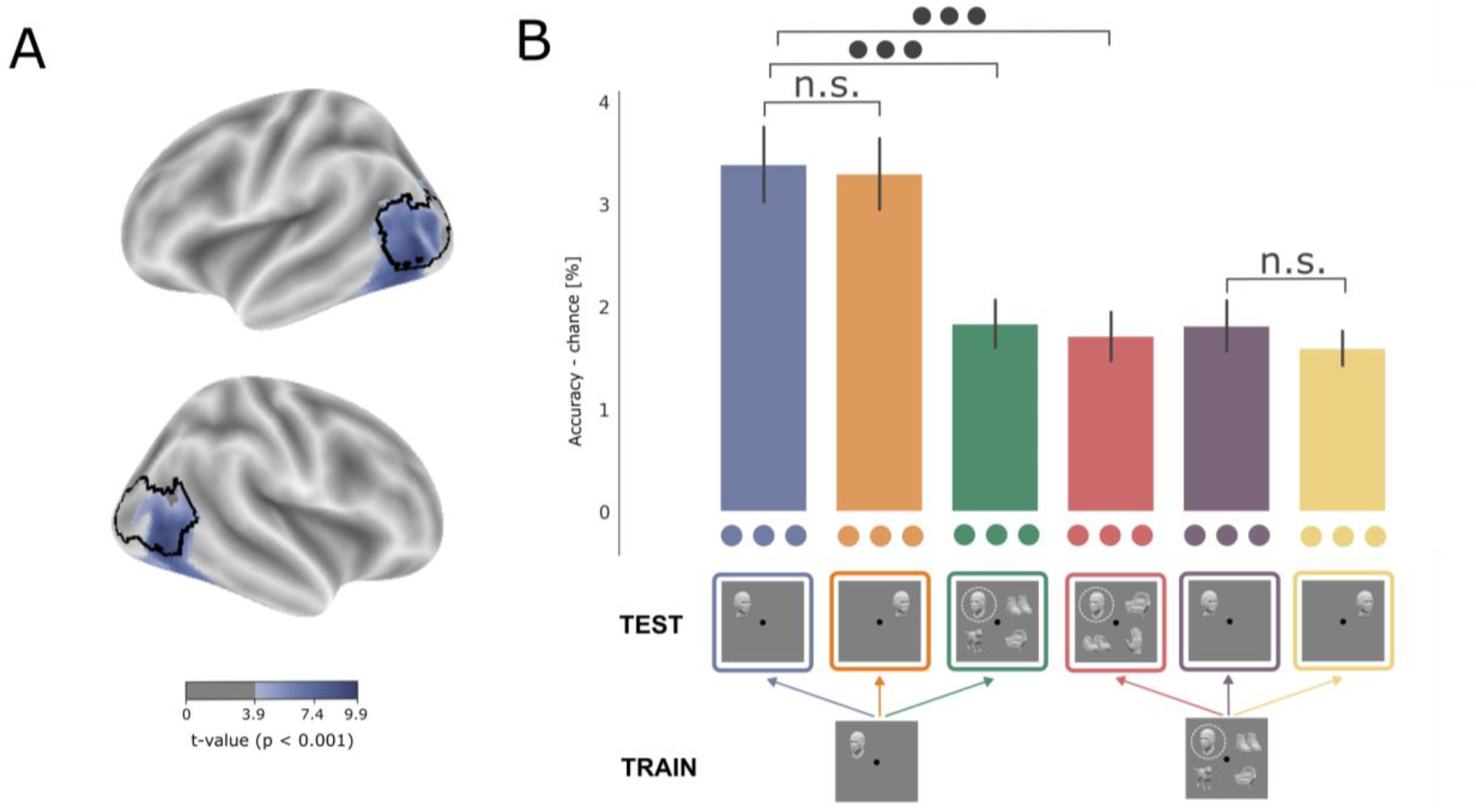
Category information in LO. **A)** ROI localization. We focused on the anatomically defined lateral occipital region (LO, outlined in black; based on probabilistic atlas by Wang et al., 2015). We confirmed that this was the locus of highest category information in a post-hoc searchlight analysis (shown in blue, see Methods). **B)** Classification logic and performance in LO. Category classifiers were trained and tested in different combinations to compare how category information generalizes across position and how sensitive it is to clutter. Classifiers were trained on trial-wise GLM parameter estimates covering the voxels in LO to distinguish between pairs of categories. Results are reported as decoding accuracy (% correct) minus chance level (50% for pairwise classification; see Methods for details). In line with previous work (MacEvoy & Epstein, 2009), results are presented as averages across object locations. Error bars show standard errors of the mean, and dots indicate significance levels (• p<0.05; •• p<0.01; ••• p<0.001; all FDR corrected).

### Decoding of single objects

As a baseline, we first trained a classifier to identify an object’s category when presented in isolation in one of the four visual field quadrants and then used this classifier to identify the category of an isolated object presented at the same location. This classifier performed with high accuracy (blue bar in **Fig. 2B**; Wilcoxon signed-rank test against 0, *p* < 0.001, *r* = 1). Next, we assessed the performance of this classifier when tested on isolated objects at a different location (i.e., in one of the other visual quadrants; orange bar in **Fig. 2B**). The classifier’s performance for this location generalization was comparably high (Wilcoxon signed-rank test against 0, *p* < 0.001, *r* = 1), with no significant difference compared to the same-location decoding (orange versus blue bars in **Fig. 2B**; Wilcoxon paired signed-rank test, *p* = 0.98, r = −0.01).

Thus, with our design we observed position-tolerant information for isolated objects. While some prior work has noted a decline in classification performance when shifting locations (e.g., Cichy et al., 2011; Kravitz et al., 2010), in our design category information for isolated objects was essentially fully position-tolerant: classifiers gained no measurable advantage from being tested at the trained position. Classifiers trained and tested in the same positions (blue bar, **Fig. 2B**) can use both position-dependent and position-tolerant representational units, whereas generalizing classifiers (orange bar, **Fig 2B**) can only rely on position-tolerant ones; the difference between the two thus reveals that very little position-dependent category information is used for classification in our design. This is an ideal foundation for testing how clutter impacts spatial tolerance. If clutter restricts LO to position-dependent information, then very little or no category information should remain decodable in clutter.

### Decoding in clutter

We then applied the classifier trained on single objects to objects at the same location but now with clutter added to the surrounding locations. This classifier also performed significantly above chance, suggesting that even in clutter, object information is still present in the LO (green bar in Fig. 2B, Wilcoxon signed-rank test against chance, p < 0.001, r = 0.99). However, we also observed a significant drop in classification performance for these cluttered object arrays compared to the isolated objects (green vs. blue bars in Fig. 2B; Wilcoxon paired signed-rank test, p < 0.001, r = 0.85), in line with previous research (Doostani et al., 2023; MacEvoy & Epstein, 2009; Reddy et al., 2009).

One possibility is that this clutter-related drop in accuracy is due to a shift in object encoding when it is surrounded by other objects compared to when it is presented alone. Various forms of spatial surround summation and modulation have been reported (Allman et al., 1985; Baeck et al., 2013; Bao & Tsao, 2018; Desimone & Duncan, 1995; Kastner et al., 1998; MacEvoy & Epstein, 2009; Miller et al., 1993). For example, invasive studies in monkey object selective regions reported averaging-like or a MAX-like behavior when multiple objects interact in the visual field (Rousselet et al., 2004; Zoccolan et al., 2005). This has also been observed in fMRI studies of human object-selective cortex (MacEvoy & Epstein, 2009; Reddy et al., 2009). If clutter changes how objects are encoded, a classifier trained on isolated objects would be poorly matched to cluttered displays, which could explain the drop in accuracy.

To test whether our drop in information after adding clutter is due to a change in the way objects are encoded in clutter, we trained similar classifiers but now in the presence of clutter (Fig. 2B, red, purple and yellow bars). If the surround really changes the encoding of object information, this classifier should be able to perform better under clutter than a classifier trained on single objects. However, when we trained classifiers to identify objects in clutter and then tested them on cluttered arrays, we did not find a recovery in classifier performance (red bar in Fig. 2B). Instead, it had a comparable drop in performance to a classifier trained on single objects (red vs. blue bars in Fig. 2B; Wilcoxon paired signed-rank test, p < 0.001, r = 0.91). This suggests that the reduction of object information in clutter is not due to a change to a “clutter-specific” encoding format. Instead, we hypothesized it may be due to a reliance on the subset of neurons with more localized, position-dependent object information.

### Spatial generalization for classifiers trained on single vs cluttered displays

We then proceeded to test whether classifiers trained in the presence of clutter indeed rely on more localized, position-dependent information. As outlined above (see **Fig. 1A**), neurons with large receptive fields are more prone to interference by the surrounding objects and thus might be less informative in the presence of clutter, whereas neurons with smaller receptive fields may be less sensitive to surround interference. A classifier that is trained only on single objects can thus utilize information from units with both small and large receptive fields. In contrast, a classifier trained in clutter should primarily rely on representational units with smaller receptive fields. If this were the case, a clutter-trained classifier is expected to generalize less well to other visual field locations than a classifier trained on isolated objects.

To test this, we evaluated clutter-trained classifiers on isolated objects, either at the trained position (purple bar, **Fig. 2B**) or at the other positions (yellow bar, **Fig 2B**). Testing on isolated objects ensures that any loss of performance at untrained positions reflects the classifier’s reliance on position-specific information rather than interference from clutter in the test data. Contrary to the prediction, both same-position and cross-position performance was well above chance (purple and yellow bars in **Fig. 2B**, respectively; Wilcoxon signed-rank test against chance, p < 0.001, r = 1 for both), and clutter-trained classifiers performed across positions as well as within (purple vs. yellow bars in **Fig. 2B**; Wilcoxon paired signed-rank test, p = 0.36, r = 0.29). This is the same pattern as for classifiers trained on isolated objects (blue and orange bars).

The single-object baseline makes this result especially informative. Because isolated objects were decoded equally well at trained and untrained positions (blue vs. orange bars), LO carried no measurable position-dependent information in our setup. Had clutter left only such position-specific information intact, there would have been little left to decode, and clutter-trained classifiers should have performed close to chance. Instead, they performed clearly above chance, both when tested in clutter (red bar, Fig. 2B) and in isolated arrays (yellow bar, Fig. 2B), and showed the same position tolerance as classifiers trained on isolated objects (yellow vs. purple bar, Fig. 2B). The information that remains decodable in clutter therefore does not appear to be restricted to the smaller sample of more position-dependent cells in LO. Instead, it seems to be the same position-tolerant information that supports the decoding of isolated objects, only reduced.

Thus, clutter does not appear to selectively degrade information encoded in position-tolerant representations in LO. This raises the question why decodable information drops from single-object to cluttered displays. A speculative, alternative possibility is that classifier performance under clutter could be disrupted by feature-based (rather than spatial) interference. It has been suggested that interference between multiple simultaneous items depends strongly on the *feature* similarity of the items (Cohen et al., 2014; Reddy & Kanwisher, 2007), and tuning sharpening has been suggested as a mechanism how attention counteracts interference between multiple stimuli (Doostani et al., 2024). Future work should address whether such feature-based interference accounts for the clutter-related drop in LO.

## Conclusion

In line with previous work, visual clutter reduced category information in LO (Doostani et al., 2023; Kastner et al., 1998; MacEvoy & Epstein, 2009; Reddy et al., 2009). However, this cost of clutter was not accompanied by a change in how category information was organized. Decoding accuracy did not improve when classifiers were trained in matching clutter conditions, and clutter-trained classifiers maintained the ability to generalize across positions. Thus, we found no evidence that clutter selectively degrades position-tolerant information presumably carried by units with large receptive fields. This challenges the idea that the visual system counteracts clutter by relying on more localized object representations. Instead, position tolerance and robustness to clutter may not be in conflict in the high-level visual cortex.

## Materials and Methods

### Participants

Nineteen healthy adults (15 females, mean age 25 years old) took part in the fMRI experiment. All reported normal or corrected-to-normal vision, and no history of neurological or psychiatric disorders. Participants gave written informed consent in accordance with the Declaration of Helsinki and received monetary compensation for their participation. The study was approved by the Ethics Committee of Humboldt-Universität zu Berlin, Department of Psychology.

### Stimulus properties and presentation

Stimulus arrays were constructed by presenting objects from the stimulus set on a gray background. Each object subtended 4° × 4° (height × width) of visual angle. Objects were centered at fixed offsets from fixation: 3° on the horizontal and 3° on the vertical axes. Objects were presented in four quadrant locations: top left, top right, bottom left, or bottom right. Luminance was equalized across the stimulus set using the SHINE toolbox (Willenbockel et al., 2010), and each object was presented on a mid-gray (RGB [0.5, 0.5, 0.5]) in isolation or in combination with other objects to create unique stimulus displays for each condition (see details below). A black fixation dot (radius 0.25°) was present continuously. Stimuli were presented on an fMRI-compatible projector with a resolution of 1920×1080 pixels and a 60Hz refresh rate, which was viewed through a mirror mounted onto the head coil at a distance of 10 cm from the participant’s eyes. Stimulus delivery and response collection were controlled with MATLAB with Psychtoolbox-3 (Brainard, 1997).

### Procedure

The experiment consisted of 3 sessions per participant, each containing 8 runs (grouped into 2 “super-runs”; see details below). In total, 24 runs (6 “super-runs”, see below) were collected across the three sessions. Each run contained 112 trials (32 single-object trials, 70 multi-object trials, 3 single-object catch-trials, and 7 multi-object catch-trials). Each run ended with an 8-15 second blank screen, and lasted a total of 7 minutes. One experimental session consisted of 56 minutes of scanning time. In session 1, a T1 anatomical image was taken at the end of the session. In sessions 2 and 3, short localizer runs were collected at the end of each session (see below). In total, each scanning session lasted around 90 minutes. Before the first scanning session, participants were familiarized with the stimulus set outside of the scanner and given instructions and 20 test trials to familiarize themselves with the task. They were given an opportunity to ask questions and clarify any uncertainties. Following the last session of the experiment, participants were debriefed about the purpose of the experiment.

Stimulus displays were presented for 250 ms per trial, followed by a variable inter-trial interval (ITI) drawn from a uniform distribution ranging between 2.5 and 3.5 s in 0.1 s increments (i.e., mean ITI was 3 s). Single-object and multi-object arrays were presented in a random order. In “catch” trials (10% of trials; see **Fig. 1C**), the stimulus was followed by a non-variable 2 second blank interval, after which the participants performed a simple 1-back task: they were shown a probe (one of the items from the stimulus set) and asked to report ‘Y’ if they saw the same object in the last presented display, and ‘N’ if they did not recall seeing the object. The probe matched the preceding stimulus array in 50% of trials; in the other 50% trials, the probe was drawn from a category that did not appear in the preceding array. The position of the ‘Y’ and ‘N’ responses on the screen (left or right) was randomized across trials. Participants gave their responses by pressing buttons with their index (left response) or middle finger (right response) on an fMRI-compatible button box with 2 buttons. The response window lasted 1.5 s, followed by visual feedback on whether the response was correct or incorrect.

A separate short localizer run was collected at the end of sessions 2 and 3. The localizer used a block design with seven blocks: faces, houses, hands, objects (chairs, shoes, cars, and plants), mixed (all eight categories), scrambled images, and baseline (blank screen). Within each block, stimulus arrays contained stimuli at all four positions, sampled with replacement; this means that four stimuli were shown simultaneously (for example, four faces, or four scrambled images), sometimes differing in orientation or exemplar, and sometimes repeating. For the block with scrambled images, each stimulus image was partitioned into 40×40 tiles, and tiles were permuted across images in the stimulus set 32 times, producing 32 scrambled images. Each block contained 20 trials lasting 16 seconds in total, with stimulus presentation lasting 0.5 s and an intertrial interval (ITI) of 0.3 s. Each localizer run comprised 7 blocks, which were repeated 2 times per run. In total, each block was repeated 4 times (2 times in session 2, and 2 times in session 3), and each localizer run lasted around 5 minutes. To maintain attention on the stimulus displays, participants performed a simple task: in 5 trials per block, one stimulus item was flipped upside-down, and participants pressed a button whenever they noticed this.

### Design and counterbalancing

We used an 8×4×2 factorial design manipulating object category (faces, hands, animals, plants, shoes, cars, chairs, houses), position (four screen locations: top-left, top-right, bottom-left, and bottom-right), and array type (single-object vs. multi-object arrays).

In single-object arrays, an object from each category was shown at each of the four positions, resulting in 32 unique object-position conditions. In multi-object arrays, for each stimulus array, 4 categories were drawn from the set of 8 categories, without repetition. This resulted in 70 unique non-ordered combinations (i.e., where position does not matter) and 1680 unique ordered combinations (i.e., where each object is selected for each position without replacement). We presented all 1680 unique arrays of 4 objects to each subject over the course of the three-session experiment. By presenting this full set of unique combinations, we ensured that each category appears equally often in each position and in combination with each other category.

To balance the 1680 multi-object arrays across runs and ensure category-position balance within runs, we wrote a custom-made optimization script in Python. First, in each run, we presented 70 multi-object trials, each being one of the 70 non-ordered unique combinations of categories (as described in the previous paragraph). Because each of the 8 categories appears in each four-item array with 50% probability (i.e., 4 out of 8 categories are presented), this results in each category being presented 35 times in each run. Second, we showed one position-permutation in each run, resulting in 24 runs. For example, one unique combination is face-house-shoes-cars, and there are 24 unique orderings across four positions (e.g., face-shoes-house-cars, cars-face-house-shoes, etc.). Third, because each category appears 35 times per run, we could not perfectly equalize the number of times it appears in each of the four positions per run (since 35 is not divisible by 4). We therefore created 6 fully-balanced “super-runs” of 4 runs, within which each category appeared exactly 35 times in each position. Satisfying these three conditions, the optimization script then ordered the position-permutations of the 70 combinations across runs and super-runs such that each run contains each of the 70 combinations, and that each category appears in each position the same number of times within and across super-runs. Subsequent decoding analyses and cross-validation were performed on the level of super-runs (i.e., leave-one-super-run-out).

The resulting matrix with balanced runs and super-runs was then used as a template to create different trials for each participant. All randomization was subject-specific and reproducible with a pseudo-random seed derived from the subject ID. First, the ordering of trials within a run, runs within each super-run, and super-runs was shuffled for each participant. Second, in each trial, one of the 4 variations of objects (2 exemplars x 2 orientations) was sampled for each category. Therefore, each participant saw the same combinations of categories, but a unique combination of exemplars and orientations. All analyses were performed only at the level of categories. Third, single-object and multi-object arrays were shown in a random order within each run, and each run contained 70 multi-object arrays (as described above) and a full set of 32 unique single-object arrays, resulting in a total of 102 trials per run. Finally, 10 trials within each run (7 multi-object trials and 3 single-object trials) were duplicated as “catch” trials and randomly intermixed with the rest of the trials. These trials were primarily used to maintain attention and vigilance during the experiment, and were excluded from analyses to avoid interference from response-related processing. This resulted in a total of 112 trials per run.

### fMRI data acquisition

Imaging data was recorded on a 3T Siemens XR Numaris MRI scanner (Siemens Healthcare, Erlangen, Germany) equipped with a 64-channel head coil and Syngo MR XA30 operating system, at the Center for Cognitive Neuroscience Berlin (Freie Universität Berlin, Germany). A high-resolution T1-weighted anatomical image was acquired at the end of the first scanning session using a 3D magnetization-prepared rapid gradient echo (MPRAGE) sequence (TR = 1900 ms, TE = 2.52 ms, TI = 900 ms, flip angle = 9°, voxel size = 1.0 × 1.0 × 1.0 mm^3^, 176 sagittal slices, field of view = 256 mm). Functional images were acquired using a T2*-weighted gradient-echo echo-planar imaging (EPI) sequence (TR = 1000 ms, TE = 30.8 ms, flip angle = 60°, multi-band acceleration factor = 6, voxel size = 2.0 × 2.0 × 2.0 mm^3^, 72 transverse slices, slice thickness = 2 mm, no gap, field of view = 220 mm). Each run consisted of 420 volumes for the main experiment and 260 volumes for the localizer run.

### fMRI preprocessing

Preprocessing was performed in SPM12 (https://www.fil.ion.ucl.ac.uk/spm/). Functional images (all 24 runs and 2 localizer runs) were first corrected for head motion using the realignment (estimate and reslice) procedure with high-quality estimation (quality = 1, separation = 2 mm, pre-smoothing FWHM = 5 mm, 7th-degree B-spline interpolation). All volumes were aligned to the first image of each run and resliced to create motion-corrected images and a mean EPI per run. The mean EPI from the first session was used as the reference for coregistration of the anatomical T1-weighted image, which was aligned to the functional space using normalized mutual information as the cost function. Coregistered anatomical images were then segmented into tissue classes (gray matter, white matter, cerebrospinal fluid, bone, soft tissue, and air/background) using SPM’s unified segmentation procedure with bias correction (bias regularization = 0.001, bias FWHM = 60 mm) and default TPM priors. All ROI analyses were performed in subject space, and apart from resampling during motion correction, functional data were not resampled again. No spatial smoothing was applied to the main experiment data. Localizer data were smoothed with a 5 mm FWHM Gaussian kernel.

### Region of interest (ROI) definition

To define LO in each participant, we used a two-step procedure combining an anatomical search space with functional voxel selection, rather than delineating LO by hand from thresholded localizer maps. This avoids subjective decisions about which activated clusters constitute the ROI (Julian et al., 2012). It also equates ROI size across participants, which matters because decoding accuracy depends on the number of voxels included.

We first defined an anatomical search space using a probabilistic atlas of human visual topography (Wang et al., 2015). Object-selective LO overlaps with regions LO1 and LO2 (Larsson & Heeger, 2006; Silson et al., 2016) and borders the hMT+ complex (Larsson & Heeger, 2006), which also shows object-selective responses (Kourtzi et al., 2002). We therefore defined a broad anatomical search space on the lateral occipital surface, combining the LO1, LO2, hMT and MST labels, thresholded at 10% and pooled across hemispheres. Early visual and ventral occipitotemporal regions were excluded. Finally, we transformed the resulting ROI into each participant’s native space using the inverse deformation fields obtained during segmentation (see Preprocessing).

Second, within this search space we selected the 350 voxels with the strongest object-selective response from the localizer run (mixed > scrambled) in each participant. Because the localizer runs were independent of the main experiment, voxel selection was independent of the decoding analyses. This yielded a subject-specific, object-selective LO ROI of equal size in every participant.

### General linear modeling (GLM)

We implemented first-level single-subject GLMs in SPM. The GLM was fit to single trials— we defined each trial as a unique regressor, and then convolved a canonical HRF with trial onset times in each run. We estimated the GLM on realigned images, and used a 128 s high-pass filter, AR(1) modeling of serial correlations, and no global normalization. Please note that single-trial estimation is known to produce noisier activation patterns and lower mean decoding accuracies than run-wise or condition-averaged GLMs, but it typically yields more stable estimates and narrower confidence intervals over cross-validation folds and participants because of reduced aggregation bias (Allefeld & Haynes, 2014; Hebart et al., 2014; Ku et al., 2008). We used these trial-wise parameter estimates as inputs to multivariate decoding.

### Multivariate category decoding

We conducted a multivariate decoding of object category within LO (see above) using functionality from the “The Decoding Toolbox” (Hebart et al., 2014). We trained linear support vector machines (SVMs) to discriminate between pairs of categories: with 8 categories, this resulted in 28 category classifiers. We trained these classifiers separately on single-object or multi-object arrays, and separately for each of the four positions, defined by the four possible stimulus locations (top-left, top-right, bottom-left, bottom-right). In single arrays, training a pairwise classifier on a particular position (e.g., top-left) involved collecting all trials where the two categories were shown in that position. In multi-object arrays, this involved collecting all trials where the two categories were shown in that position, with an important additional constraint that we excluded any trials where both decoded categories appeared in the same multi-object array. For example, when training a top-left face-vs-house classifier, we included only the trials where a face appeared in top-left, and a house did not appear in the other three positions, and conversely, only the trials where a house appeared in top-left and a face did not appear in any other positions. In total, this resulted in 224 classifiers (28 pairs, four positions, and 2 array conditions).

We then tested the classifiers in different combinations, depending on the analysis. For position-dependent analysis, we trained and tested the classifier on the same position. For position generalization analysis (position-tolerant analysis), we trained the classifier on one position and tested its performance separately on each of the three other positions. For clutter generalization analyses, we trained the classifier on one position in single-object arrays and tested its performance in the same position in multi-object arrays, and vice versa. Finally, for position generalization of clutter-trained classifiers, we trained the classifier on one position of multi-object arrays and tested its performance separately on each of the three other positions in single-object arrays.

For cross-validation, runs were grouped into 6 super-runs, each containing 4 runs (see Design and counterbalancing). We performed six-fold leave-one-super-run-out cross-validation, training on trials from 5 super-runs, and testing on the left-out 1 super-run. For single-object trials, this meant that each classifier was trained on 40 trials (20 for each label) and tested on 8 trials (4 for each label). In order to match sample sizes, for multi-object trials, we subsampled the number of trials to match the number of available single-trials by randomly selecting 40 trials for training (20 for each label) and 8 trials for testing (4 for each label). The same cross-validation scheme was used in position-dependent and generalization analyses—even when we trained and tested classifiers on different data, we still only used ⅚ of the available data for training. This ensured comparability across train-test regimes.

As classification output, we computed classification accuracy above chance level (50%). For each participant, we averaged classification accuracies across category classifiers. For position-dependent classifiers, this directly resulted in one estimate per participant. For position-generalization classifiers (i.e., trained on one position and tested separately on three other positions), we additionally averaged classifier performance across the three positions, resulting in one mean estimate per participant.

### Searchlight analysis

To confirm that our ROI captured the main locus of category information, we performed a post-hoc searchlight analysis using The Decoding Toolbox (Hebart et al., 2014). For computational efficiency, this analysis differed from the ROI analysis in that it used parameter estimates from a condition-wise first-level GLM (one regressor per category × position × array type in each run) instead of single-trial estimates, and that it used a single eight-way category classification instead of 28 pairwise classifiers. Only single-object arrays were included. In each participant, a sphere with a radius of 4 voxels (8 mm) was moved across all gray-matter voxels, as defined by SPM segmentation of the participant’s anatomical image. Within each sphere, a linear support vector machine was trained to discriminate all eight object categories, trained and tested within the same position, and using the same six-fold leave-one-super-run-out cross-validation as in the ROI analysis. Individual accuracy maps were then normalized to MNI space using the forward deformation fields obtained during segmentation and resampled to 2 mm isotropic voxels with 4th-degree B-spline interpolation. We tested accuracy minus chance against zero at the group level with voxel-wise one-sample t-tests across participants and thresholded the resulting t-map at p < 0.001 (two-sided, uncorrected). Voxels exceeding this threshold are shown in blue in **Fig. 2A**.

### Statistical analysis

Pairwise comparisons of classifier performance, as well as comparisons of decoding accuracies against chance, were performed using the Wilcoxon signed-rank test, appropriate for non-n?ormally distributed, repeated-measures data. Zero differences were excluded following the standard Wilcoxon procedure. In addition to reporting the p-value, we quantified effect size using the rank-biserial correlation, calculated as the relative balance of positive and negative signed ranks. This measure ranges from −1 to 1 and provides an interpretable index of the magnitude and direction of the paired difference. When multiple pairwise Wilcoxon comparisons were performed within the same family of tests, we applied false discovery rate (FDR) correction to control for the expected proportion of false positives. FDR correction used the Benjamini–Hochberg procedure (Benjamini & Hochberg, 1995) at *q* = 0.05. All reported significance levels in the Results reflect FDR-adjusted *p*-values unless stated otherwise.

## Acknowledgements

We thank Karolis Degutis and Rotem Krispil for helpful discussions of the results reported here, and we thank the staff of the Center for Cognitive Neuroscience Berlin (Freie Universität Berlin) for support with fMRI data acquisition. K.M. was funded by the Max Planck Society and BMBF (as part of the Max Planck School of Cognition). J.D.H. was supported by the Deutsche Forschungsgemeinschaft (DFG, Exzellenzcluster Science of Intelligence); SFB 940 “Volition and Cognitive Control”; and SFB-TRR 295 “Retuning dynamic motor network disorders using neuromodulation”.

